# A Novel Lossless Encoding Algorithm for Data Compression - Genomics Data as an Exemplar

**DOI:** 10.1101/2020.08.24.264366

**Authors:** Anas Al-okaily, Abdelghani Tbakhi

**Affiliations:** Department of Cell Therapy and Applied Genomics, King Hussein Cancer Center, Amman, Jordan; Department of Pathology and Molecular Medicine, McMaster University, Ontario, Canada

**Author notes:** Email addresses:* (Anas Al-okaily), (Abdelghani Tbakhi).

**Keywords:** DNA compression, data encoding, Huffman encoding

## Abstract

Data compression is a challenging and increasingly important problem. As the amount of data generated daily continues to increase, efficient transmission and storage has never been more critical. In this study, a novel encoding algorithm is proposed, motivated by the compression of DNA data and associated characteristics. The proposed algorithm follows a divide-and-conquer approach by scanning the whole genome, classifying subsequences based on similarity patterns, and binning similar subsequences together. The data are then compressed in each bin independently. This approach is different than the currently known approaches: entropy, dictionary, predictive, or transform based methods. Proof-of-concept performance was evaluated using a benchmark dataset with seventeen genomes ranging in size from kilobytes to gigabytes. The results showed considerable improvement in the compression of each genome, preserving several megabytes compared with state-of-art tools. Moreover, the algorithm can be applied to the compression of other data types include mainly text, numbers, images, audio, and video which are being generated daily and unprecedentedly in massive volumes.

## 1. Introduction

The importance of data compression, a fundamental problem in computer science, information theory, and coding theory, continues to increase as global data quantities expand rapidly. The primary goal of compression is to reduce the size of data for subsequent storage or transmission. There are two common types of compression algorithms: lossless and lossy. Lossless algorithms guarantee exact restoration of the original data, while lossy algorithms do not. Such losses are caused, for instance, by the exclusion of unnecessary information, such as metadata in video or audio that will not be observed by users.

Data exist in different formats including text, numbers, images, audio, and video. Several coding algorithms and corresponding variants have been developed for textual data, the primary focus of this paper. This includes the Huffman Huffman et al. (1952), Shannon Shannon (2001), Shannon-Fano Fano (1949), Shannon-Fano-Elias Cover (1999), Lempel-Ziv (LZ77) Ziv and Lempel (1977), the Burrows-Wheeler transform Burrows and Wheeler (1994), and Tunstall Tunstall (1967) algorithms. The Huffman algorithm includes several variants: minimum variance Huffman, canonical Huffman, length-limited Huffman, non-binary Huffman, adaptive Huffman, Faller-Gallager-Knuth (an adaptive Huffman) Knuth (1985), and Vitter (an adaptive Huffman) Vitter (1987). The LZ algorithm also includes several variants, such as LZ78 Ziv and Lempel (1978), Lempel-Ziv-Welch (LZW) Welch (1984), Lempel-Ziv-Stac (LZS) Friend (2004), Lempel-Ziv-Oberhumer (LZO) Oberhumer (2008), Lempel-Ziv-Storer-Szymanski (LZSS) Storer and Szymanski (1982), Lempel–Ziv-Ross-Williams (LZRW) Williams (1991), and the Lempel–Ziv–Markov chain algorithm (LZMA) Ranganathan and Henriques (1993). Additional techniques involve arithmetic encoding Langdon (1984), range encoding Martín (1979), move-to-front encoding (also referred as symbol ranking encoding) Ryabko (1980); Bentley et al. (1986), run-length encoding Capon (1959), delta encoding, unary encoding, context tree weighting encoding Willems et al. (1995), prediction by partial matching Cleary and Witten (1984), context mixing Mahoney (2005), asymmetric numeral systems (also called asymmetric binary coding) Duda (2013), length index preserving transform Awan and Mukherjee (2001), and dynamic Markov encoding Cormack and Horspool (1987).

Compression algorithms can be classified based on the methodology used in the algorithm, such as entropy, dictionary, predictive, and transform based methods. These methods have been described extensively in several recent studies Gopinath and Ravisankar (2020); Kavitha (2016); Uthayakumar et al. (2018); Kodituwakku and Amarasinghe (2010), however, a brief description for each method is provided in the Supplemental Methods.

Genomics (DNA/RNA) data is a type of textual information with several unique characteristics. First, the alphabet consists only of A, C, G, and T characters representing the four nucleotides: adenine, cytosine, guanine, and thymine, respectively. Second, DNA data contain repeat sequences and palindromes. Third, the size of genomics data can be very large, relative to most media files. The human genome, for instance, consists of more than three billion nucleotides (specifically 3,272,116,950 bases (https://www.ncbi.nlm.nih.gov/grc/human/data?asm=GRCh38.p13) requiring more than three gigabytes of storage). As such, sequencing genomic data (especially for humans) is currently being performed for research and diagnostic purposes in daily basis. In addition, this sequencing is typically conducted with high depth (30-100x) to sequence the same region several times in order to make reading the DNA regions more accurate. This generates massive quantities of data on a daily basis. For example, the number of bases sequenced from December 1982 through December 2022 was 19,086,596,616,569 (https://www.ncbi.nlm.nih.gov/genbank/statistics/). Several algorithms have been developed to compress these data, which can be divided into vertical and horizontal techniques Grumbach and Tahi (1994). Vertical mode algorithms utilize a reference genome/source, while horizontal mode algorithms are reference-free.

Genomic data are stored in different formats, including FASTA Lipman and Pearson (1985), FASTQ Cock et al. (2010), and SAM Li et al. (2009), with FASTA being the most common and also the primary focus of this paper. Several comparative studies for compressing FASTA files have been published in recent years Kryukov et al. (2020); Hosseini et al. (2016); Mansouri et al. (2020); Bakr et al. (2013); Jahaan et al. (2017). Genomic sequences typically consist of millions or billions of sequenced reads, with lengths in the hundreds, stored with the quality of each base in a primarily FASTQ format. A common DNA data processing strategy involves aligning the sequenced reads with a reference genome. The output is the reads themselves, with their base qualities and alignment results for each read, stored in a SAM format. Surveys of compression tools for SAM and FASTQ data are available in the literature Bonfield and Mahoney (2013); Hosseini et al. (2016).

The small alphabet found in DNA data simplifies the compression process. However, the problem remains challenging due to the discrete, uniform distribution (frequencies) of bases in DNA data. Efficient compression relies mainly on repetitiveness in the data and encoding as few characters/words as possible, since encoding more characters costs more bits-per-character. If the characters are uniformly distributed in the text, their repetitions will also be distributed uniformly and encoding only a fraction of them (to decrease the bits-per-character) will lead to low compression outcomes. The application of Huffman encoding, for instance, produces 2-bit assignments for each base. The algorithm will then produce an inefficient/suboptimal compression result that does not utilize repetitions found in the DNA data. Motivated by this issue, we introduce in this work a lossless and reference-free encoding algorithm.

## 2. Methods

The following observations can be inferred from a careful analysis of DNA. First, many regional (local) sub-sequences (assume a length of 100 bases) contain non-uniform or skewed distributions. Second, similar sub-sequences (which provide better compression results if encoded together) are often distant from each other. This distance is typically longer than the length of sliding windows (usually in kilobases/kilobytes) commonly used in algorithms such as LZ or far apart from previous sequence/symbol used in prediction models such as context weighting tree, predictive partial matching, or dynamic Markov compression. Even if these sequences are located within the same sliding window or previous sequences/symbols, they are often sufficiently distant from each other, which leads to inefficient compression and encoding. These two observations were the motivation behind the design of the following encoding algorithm.

### 2.1. Compression algorithm

Given a string *T* of length *t*, containing characters from a fixed alphabet of length Σ, and a window-length *w*, the description of the proposed algorithm is stated in Algorithm 1.

#### Algorithm 1

Compression algorithm

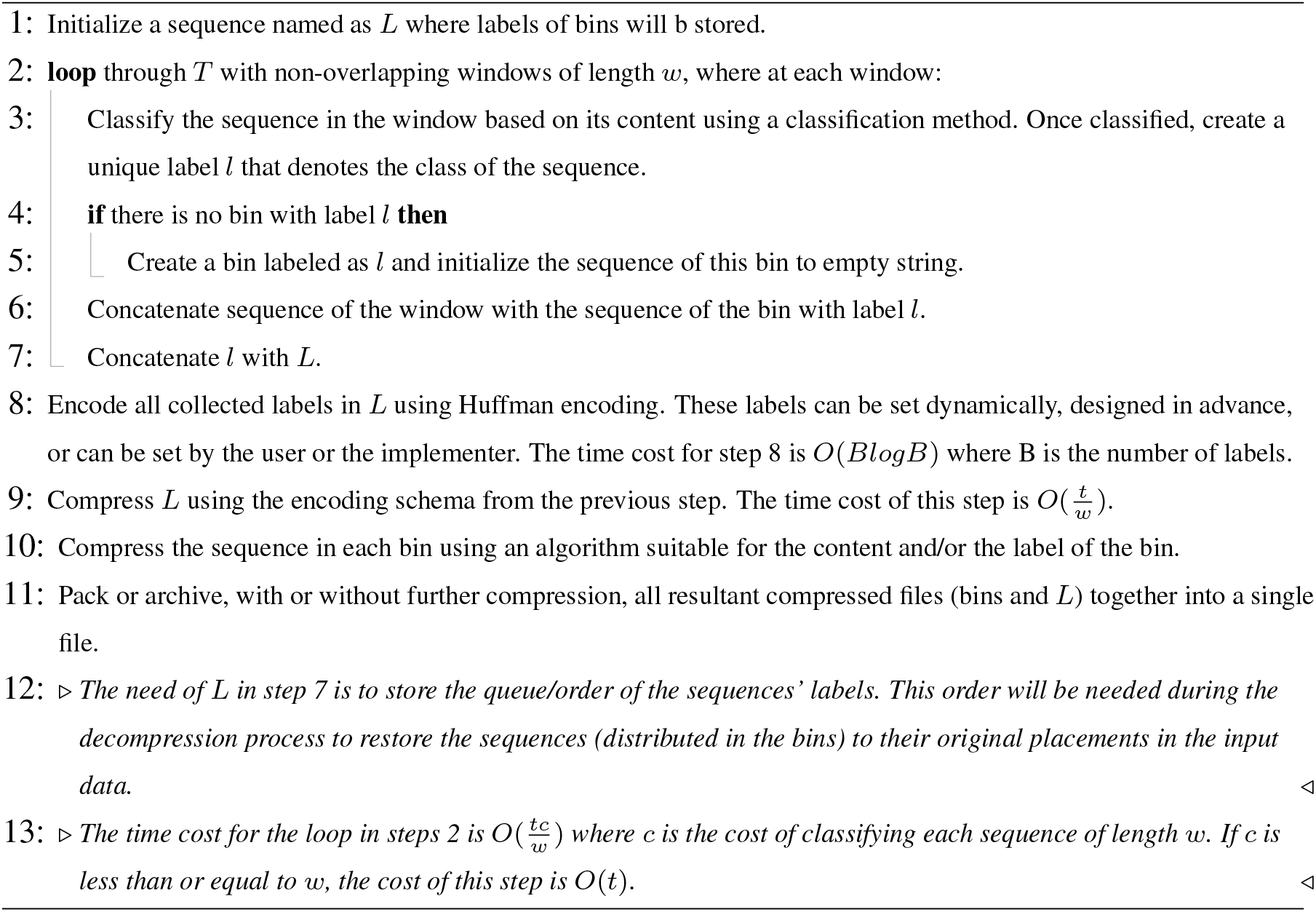

Note the value of *w* can be fixed or variable. If *w* is a variable, the window is extended character by character until the label of the sequence in the window matches one of the bin labels. The length of the window must then be added to *L* (after the string representing the label for each window) or placed in a separate stream that stores lengths and their order. Lastly, lengths can be encoded using a universal code for integers (such as Levenshtein coding (http://www.compression.ru/download/articles/int/levenstein_1968_on_the_redundancy_and_delay.pdf), Elias coding Elias (1975) (delta, gamma, or omega), exponential-Golomb code Salomon (2004), Fibonacci code Fraenkel and Kleinb (1996), Stout code Stout (1980)) or using a suitable encoding algorithm such as unary, binary, or Huffman.

### 2.2. Decompression algorithm

Decompression algorithm is the inverse of compression algorithm and is described as stated in Algorithm 2.

#### Algorithm 2

Decompression algorithm

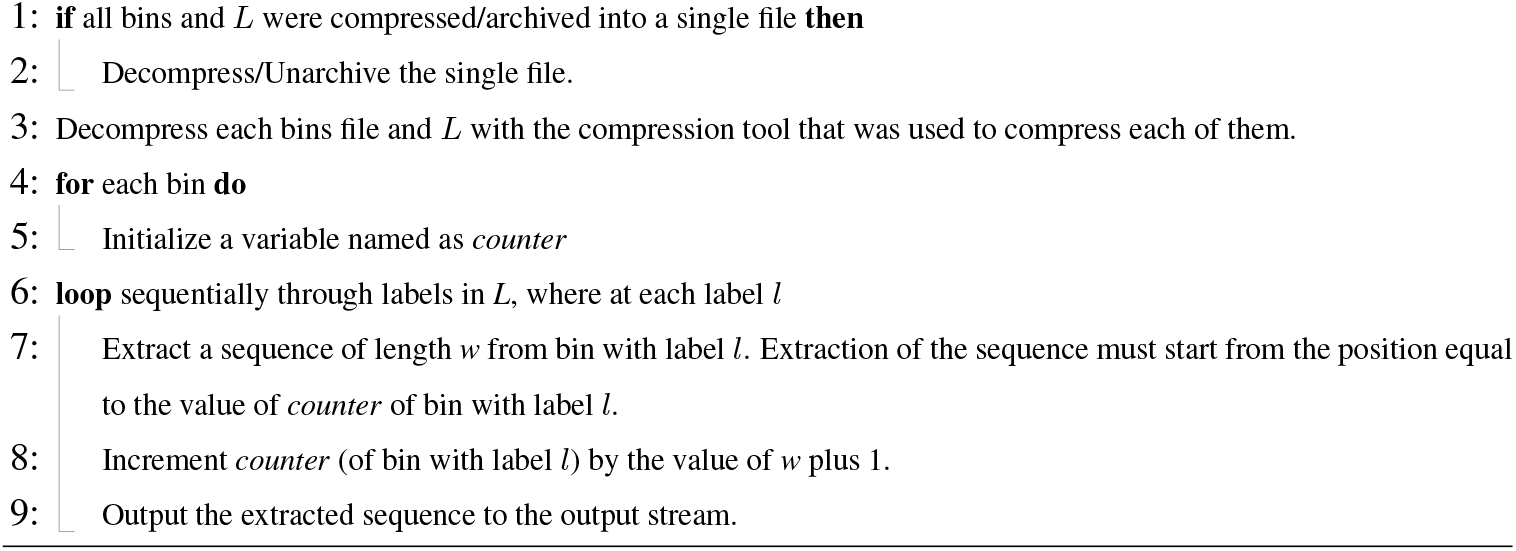

The time and memory cost of decompression is linear.

This algorithm can be applied not only to DNA or textual data, but to archiving processes and other data types namely numbers, images (binning for instance similar pixels instead of similar subsequences as in text), audio (binning for instance similar subbands/frequency-ranges), and video (binning for instance similar images)). Sub-binning or nested-binning processes can also be applied.

This design facilitates organizing and sorting the input data using a divide-and-conquer method by creating bins for similar data and encodes/compresses data in the same bin that are better of compressed together, to achieve better compression results with a minor increase in time costs. In the case of more random/divergent data, which is common, this is done to avoid relying on a single encoding or compression technique (as in entropy methods), being dependent on the randomness of previous sequences (as in prediction methods), requiring construction of dictionaries dependent on the randomness of previous sequences (as in dictionary methods), or potentially introducing inefficient transformation due to the randomness of the data (as in transform methods). In contrast, the proposed algorithm divides the data into groups of similar segments, regardless of their position in the original sequence, which decreases the randomness and contributes in organizing the input data to ultimately handling the compression process more efficiently.

Note that the continued application of sub-binning processes will eventually reduce the randomness/divergence of the data and improve the compression results, by obtaining data that are optimal or suboptimal for compression. These processes will require additional time costs, but these costs will still be practical at low sub-binning depth and feasible at high sub-binning depths, especially for small data or the compression of large data for archiving. Therefore, sub-binning will eventually provide more control, organization, and possibly a deterministic solution to encoding and compression problems. Further analysis and investigations are also provided in the Supplemental_Methods.

This encoding algorithm is named in honor of Pramod Srivastava (Professor in the Department of Immunology, University of Connecticut School of Medicine) who was an advisor to the first author. As such, it is named the Okaily-Srivastava-Tbakhi (*OST*) algorithm.

### 2.3. OST-DNA

This is the first implementation of the OST algorithm which accepts DNA data as input which can be denoted as OST-DNA. Note that the aim of this implementation is to proof-the-concept of the proposed algorithm. Academic and commercial versions and after careful sophistication and customized methods will be sought in the near future. Bin labels are computed using a Huffman tree encoding strategy. For example, if the Huffman tree for a subsequence produces the following encoding schema: G:0, A:10, T:110, and C:111, then the label will be GATC 1233 label (1 indicates G is encoded using 1 bit, A using 2 bits, and so on). The number of bits used to encode a character gives a general indication of its frequency compared to the other characters. The number of bins can be reduced by decreasing the label length as follows. To produce a label length of 1, we used the first base of the Huffman tree and its bit length. As such, the above Huffman encoding schema will be represented by G 1. If the bin label length is 2, then the label will be GA 12. This clearly decreases the number of labels, but at the cost of decreasing the similarity among sequences in the same bin therefore their compression. Note that this classification method (bin labeling) is suitable for DNA data as its alphabet is small. For data with larger alphabets, same or other classification methods might be sought such as binning sequences that contain some letter/s most frequently.

As the windows do not overlap, each base in *T* will be read in *O*(1) time. The cost of constructing a Huffman tree for a subsequence is then *O*(Σ*log*Σ), requiring the construction of 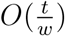 Huffman trees for all subsequences in *T*. The total cost hence is 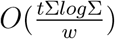. In order to allow for the acquisition of non-uniform distributions for the characters in Σ (the pigeonhole principal), the value of *w* must be larger than that of Σ*log*Σ, noting that Σ is a constant. As such, the total cost the compression process of OST-DNA is *O*(*t*).

Since the value of *w* is fixed in this version of OST-DNA, Huffman trees are constructed once for each window sequence. In the case of a variable *w* where the window will be extended until the sequence label matches a bin label, it is not efficient to calculate Huffman trees for the entire sequence at every extension, hence adaptive Huffman trees can be applied instead. Generally, the compressed bin files and *L* can be collected into a single file using an archiver which could perform further compression. However, this process was omitted in this study to demonstrate the efficiency of the OST algorithm without any further compression may be produced by the archiving process.

## 3. Results

We implemented OST-DNA using the python language. We used the same dataset applied to another benchmark Kryukov et al. (2020) in order to test and compare the compression results from OST-DNA with the tools listed in Table S1 in Supplemental Methods. The dataset consists of seventeen genomes, as shown in Tables S2 and S3 in Supplemental_Methods.

The following preprocessing steps were applied to each tested genome. All new lines, header lines, lowercase bases, and bases not identified as A, C, G, T, or N, were removed. This produced a one-line sequence for each genome, containing only the four capitalized DNA bases and the letter “N”. Character “N” represents uncalled/undetermined base during sequencing or assembling process. Assembled genomes may contain also bases with lowercase that represent softmasked sequences. In compression process, these sequences are converted to capital case with recording their start/end coordinates so that during decompressing process their original case is restored. As the tested genomes contain low number of soft-masked sequences and as this study is using genomics data for testing purposes, these sequences were just removed from the genomes. The python script used to perform these preprocessing steps and the size of each genome, before and after applying the script, are provided in Table S4 in Supplemental_Methods. The size of one-line genomes ranged from 50KB to 13GB with a total size of 16,773.88MB.

Compression ratio was the primary metric used for evaluating the performance of the proposed algorithm. It is equal to the size of the compressed file divided by the size of the uncompressed (original) file. The original files in this study are the one-line genome files. Other metrics include compression time (seconds), decompression time (seconds), compression speed (the size of the uncompressed file in MB divided by the compression time in seconds-MB/s), and the decompression speed (the size of the uncompressed file in MB divided by the decompression time in seconds-MB/s).

The following tools were selected as they are common tools for compressing textual data and implementing one or more compression algorithms available in the literature. This is meant to test compressing the resultant bins using all available encoding algorithms. The tools namely are: bcm, blzpack, brotli, bsc, bzip2, cmix, compress, freeze, gzip, hook, Huffman-codec, lizard, lrzip, lz4, lzb, lzfse, lzip, lzop, lzturbo, Nakamichi, ppm, qzip, rans static, rzip, snzip, srank, xz, zlib, zip, zpipe, and zstd. Description of each tool is provided in Table S1 in Supplemental_Methods.

Each tool was applied to each one-line genome. The compression and decompression commands used to run each tool are provided in Table S5 in Supplemental_Methods. The default options for each tool were used to compress the one-line genomes. The same commands (default options) were used to compress the bins. No parallel processing is applied. As some tools apply parallel processing by default as a result compression/decompression time will be affected to the favor of these tools; if the default options of a tool is to apply parallel processing, the options were modified to be single/sequential processing. The cumulative compression results (for all one-line genomes) are provided in Table 1, while the compression results for each one-line genome are listed in S6_Table. The most efficient tools in terms of compression ratio were lrzip (saved 14,317.40 MB out of 16,773.88 MB), brotli (13,958.42 MB), lzip (13,916.39 MB), xz (13,915.92 MB), bsc (13,391.09 MB), and bcm (13,314.83 MB). In addition, comparing the results of the commonly used tools (bzip2 and gzip) indicated bzip2 was better, saving 12,601.12 MB.

**Table 1:**
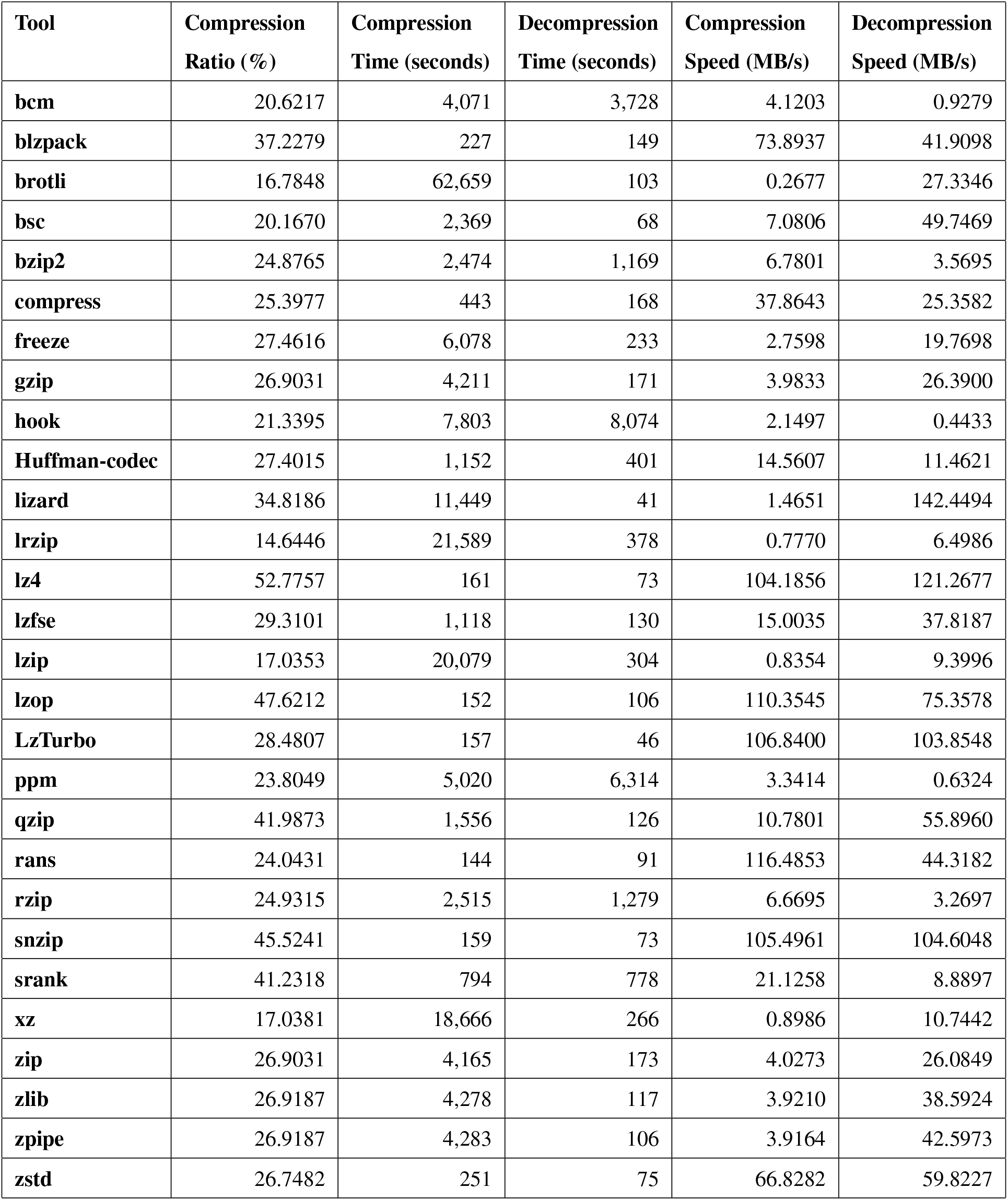
Compression performance for each common tool cumulatively over all tested genomes. The size of all genomes (in one-line format) is 16,773.88 MB. The tools cmix, lzb, and Nakamichi could not compress large genomes in reasonable time so their cumulative performance could not be presented.

Seven versions of OST-DNA were implemented. In each version, one of the seven most efficient tools (bcm, brotli, bsc, lrzip, lzip, xz, and bzip2) is used to compress the bins generated by the OST-DNA algorithm. The command used by each tool to compress the one-line genomes is the same used to compress the bins. Each of these seven versions were run over each window lengths of 25, 50, 100, 125, 150, 250, 500, 750, 1,000, 2,500, 5,000, and 10,000 and across label lengths of 1, 2, 3, 4, and 5. The best result in terms of compression ratio over all pairs of window lengths and label lengths and cumulatively (over all 17 one-line genomes) achieved by each OST-DNA version is shown in Table 2.

**Table 2:**
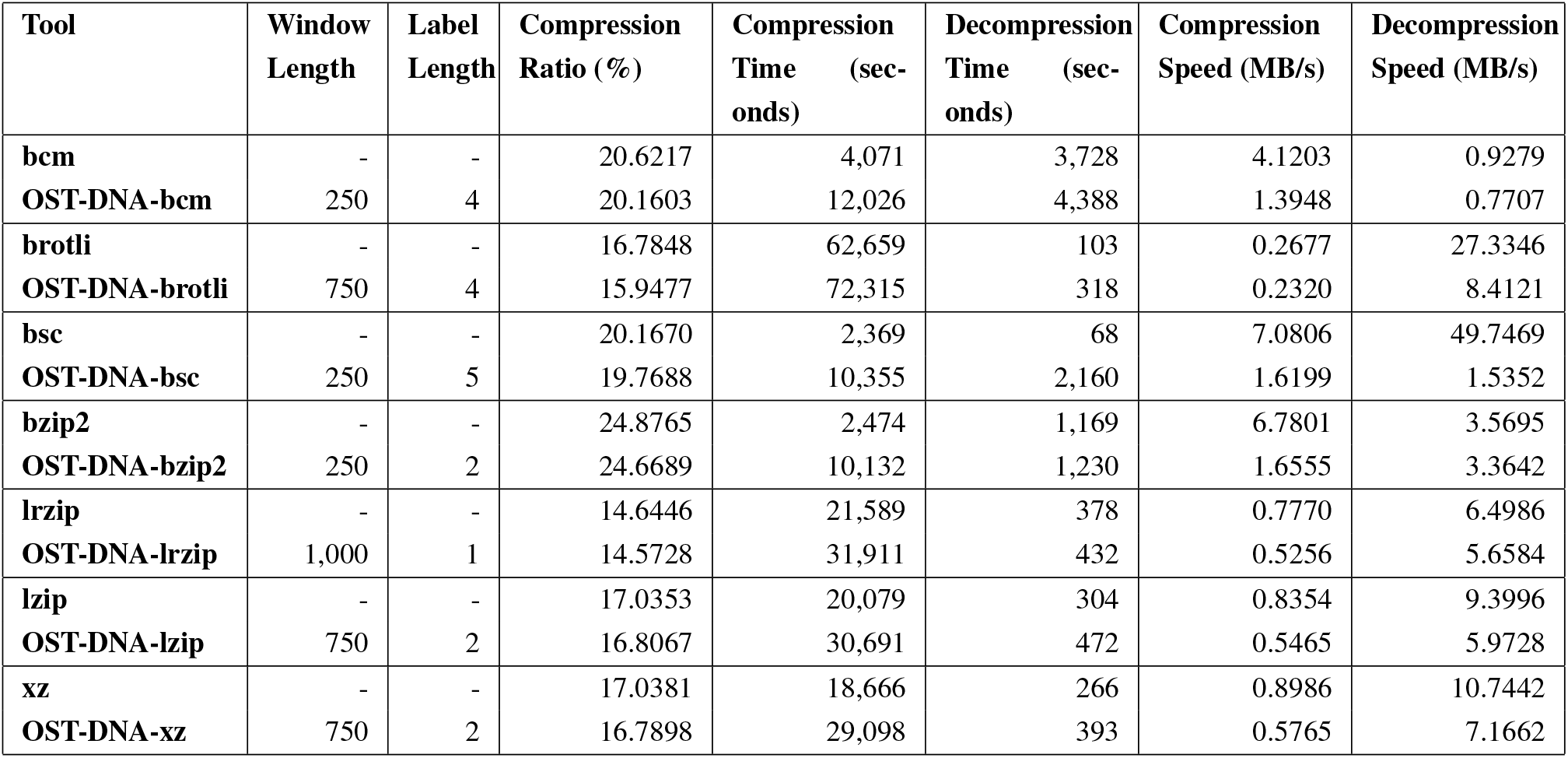
Compression performance for best window-length and label-length of each of the seven OST-DNA versions cumulatively over all tested genomes.

A comparison of the results produced by each OST-DNA tool (i.e., bcm, brotli, bsc, bzip2, lrzip, lzip, and xz) indicated OST-DNA-bcm saved an additional 77.38 MB compared to bcm, OST-DNA-brotli: 140.41 MB, OST-DNA-bsc: 66.79 MB, OST-DNA-bzip2: 34.83 MB, OST-DNA-lrzip: 12.05 MB, OST-DNA-lzip: 38.34 MB, and OST-DNA-xz: 41.65 MB. This demonstrates the proposed algorithm can improve compression results compared to the individual corresponding tools.

The best tool in terms of compression ratio was lrzip, yet OST-DNA-lrzip saved an additional 12.05 MB more than lrzip. In terms of compression time, bsc was the fastest tool. OST-DNA-bsc could save an additional 66.79 MB more than bsc with a practical increase in the compression/decompression times (hence corresponding decrease in compression and decompression speeds). These increases are a result of the time needed for classifying and binning sequences during compression, as well as the need to collect and restore the original genome during decompression. However, they can be decreased significantly as follows. First, the OST-DNA script was not optimized for implementation as it is intended in this study to provide proof of concept. Additional improvements to the script can reduce both the compression and decompression times by increasing the corresponding speeds. In addition, parallel processing, which could further reduce run-time, was not applied during the binning, compression, or decompression steps of OST-DNA. Finally, fewer bins would lead to faster sequence labeling and longer window lengths could speed up both compression and decompression, with a trade-off in the compression ratio.

The compression results for OST-DNA using each of the seven tools for each one-line genome were also better than the results using the corresponding standalone tool. This can be found by comparing the compression results using each OST-DNA version with each window length and each label length for the one-line genomes, as shown in S7_Table. The compression results for the standalone tools are provided in Table 1.

By analyzing the compression results for all OST-DNA versions, using different sequence label lengths and classification methods (i.e., Huffman tree encoding schema), we found the most efficient results correlated with a window length of 250 to 1,000 bases. This is reasonable, as lengths shorter or longer than this will yield a uniform distribution of bases in the sequence. However, dynamic window lengths can be more practical and feasible given the additional costs for encoding the lengths. We found efficient label lengths to be 2 and 4. This is reasonable as increased label lengths produce more bins and more similarities among sequences in the same bin. Compression is more efficient when sequences in a bin are more similar. Additional classification methods can be used to label the bins and improve compression further. S8_Table shows compression results for each version of OST-DNA for each window and each label length cumulatively applied to all one-line genomes. Further analysis at the bin level, rather than the genome level, is provided in S9_Table.

Compression results produced by applying each OST-DNA version to each bin, using the same window and label lengths but with different genomes, were considerably different (see S10_Table). This was not the case for bins produced using the same label length and same genome, but with different window lengths (see S11_Table). This means that sequences with the same label but from different genomes differed significantly (even though their labels were the same). This observation suggests the need to find a set of labels or labeling steps that could compress sequences from any source (genome) with similar efficiency, to improve the compression results further. In other words, sequences that share a label from this set would be compressed at a similar rate, regardless of the source (genome) from which they were derived. This set of labels could also be used better archival of multiple genomes.

The current version of OST-DNA compresses each bin using a specific tool. However, this is not optimal. Finding a tool that optimally compresses each bin, or a novel algorithm that is customized for efficient compression based on the bin content or label, could further improve the overall performance.

## 4. Code availability

Source code of the seven OST-DNA tools are available at https://github.com/aalokaily/OST.

