## Supplemental_Methods for "A Novel Lossless Encoding Algorithm for Data Compression - Genomics Data as an Exemplar"

### Material

In order to benchmark OST-DNA and the common compression tools, we used the tools provided in Table S1 with their descriptions. In addition, we used the genomes in Tables S2 and S3.

### Methods

As discussed in the main text, all newlines, header lines, bases that are not A, C, G, T, or N were removed from the tested genomes to obtain a one-line sequence format for the each genome. The python script that was used for this purpose is stated below.

```
import sys
import os
import fileinput

fasta_file = sys.argv[1]
out_file =  open(os.path.basename(fasta_file).rsplit(".",1)[0] + ".one_line", 'wa')

# get all variants from each file without intersection
for line in fileinput.input(fasta_file):
    if line[0] == ">":
        pass
    else:
        for c in line:
            if c in ["A", "C", "G", "T", "N"]:
                out_file.write(c)
```

The size of each genomes before and after applying the python script is stated in Table S4.

Table S5 stated the commands used to compress one-line genomes for each tool. The same command which was used for bcm, brotli, bsc, bzip2, lrzip, lzip, and xz was also used to compress the bins in OST-DNA-bcm, OST-DNA-brotli, OST-DNA-bsc, OST-DNA-bzip2, OST-DNA-lrzip, OST-DNA-lzip, and OST-DNA-xz; respectively.

For unknown reason and given that the latest version of lrzip is v0.631, none of the versions later than v0.50 worked properly for the proposed algorithm, namely, in the step of compressing bins. So, v0.50 the version was used in this work.

The coding language which was used to implement OST-DNA versions is python 2.7.

The operating system and machine used for running and testing is as follows.

**$ cat /etc/os-release**

```
NAME="CentOS Linux"
VERSION="7 (Core)"
ID="centos"
ID_LIKE="rhel fedora"
VERSION_ID="7"
PRETTY_NAME="CentOS Linux 7 (Core)"
ANSI_COLOR="0;31"
CPE_NAME="cpe:/o:centos:centos:7"
HOME_URL="https://www.centos.org/"
BUG_REPORT_URL="https://bugs.centos.org/"
CENTOS_MANTISBT_PROJECT="CentOS-7"
CENTOS_MANTISBT_PROJECT_VERSION="7"
REDHAT_SUPPORT_PRODUCT="centos"
REDHAT_SUPPORT_PRODUCT_VERSION="7"
```

**$ lscpu**

```
Architecture:          x86_64
CPU op-mode(s):        32-bit, 64-bit
Byte Order:            Little Endian
CPU(s):                64
On-line CPU(s) list:   0-63
Thread(s) per core:    2
Core(s) per socket:    16
Socket(s):             2
NUMA node(s):          2
Vendor ID:             GenuineIntel
CPU family:            6
Model:                 79
Model name:            Intel(R) Xeon(R) CPU E5-2683 v4 @ 2.10GHz
```

Stepping: 1
CPU MHz: 1202.142
CPU max MHz: 3000.0000
CPU min MHz: 1200.0000
BogoMIPS: 4199.62
Virtualization: VT-x
L1d cache: 32K
L1i cache: 32K
L2 cache: 256K
L3 cache: 40960K

| **Tool** | **Algorithm** | **Version** | **Year** | **Source Code** |
|---|---|---|---|---|
| bcm | Burrows-Wheeler transform | v1.51 | 2020 | https://sourceforge.net/projects/bcm/ |
| blzpack | brieflz library (https://github.com/jibsen/brieflz) which implements LZ style compression | v1.3.0 | 2020 | https://github.com/jibsen/brieflz/tree/master/example |
| brotli | Google tool uses LZ77, Huffman encoding, and context modelling | v1.0.7 | 2018 | https://github.com/google/brotli |
| bsc | Block sorting | v3.1.0 | 2016 | https://github.com/IlyaGrebnov/libbsc |
| bzip2 | Burrows-Wheeler transform | v1.0.8 | 2019 | http://www.bzip.org/ |
| cmix | Model prediction and Context mixing | v18 | 2019 | https://github.com/byronknoll/cmix |
| compress | LZW algorithm | v4.2.4 | 2011 | https://github.com/vapier/ncompress |
| freeze | LZSS and Huffman encoding | v2.5.0 | 1993 | https://centos.pkgs.org/7/epel-x86_64/freeze-2.5.0-16.el7.x86_64.rpm.html |
| gzip | LZSS and Huffman coding | v1.10 | 2018 | https://ftp.gnu.org/gnu/gzip/ |
| hook | Dynamic Markov encoding | v0.8e | 2007 | http://cs.fit.edu/~mmahoney/compression/hook.zip |
| Huffman-codec | Huffman encoding | v1.0 | 2020 | https://github.com/Bestoa/huffman-codec |
| lizard | LZ77 and Huffman encoding | v1.0 | 2019 | https://github.com/inikep/lizard |
| lrzip | Extension of rzip but replacing bzip2 by LZMA, LZO, or no second-stage | v0.50 | 2010 | http://ck.kolivas.org/apps/lrzip/ |
| lz4 | LZ77 family of byte-oriented compression schemes | v1.9.2 | 2019 | https://github.com/lz4/lz4 |
| lzb | Hexadecimal and base64 encoding/decoding | v1.0 | 2019 | https://github.com/faragon/lzb |
| lzfse | Apple tool that applies LZ style compression using finite state entropy | v1.0 | 2017 | https://developer.apple.com/documentation/comp |

| | | | | |
|---|---|---|---|---|
| | | | | ression/algorithm/lzfse |
| lzip | LZMA | v1.19 | 2017 | https://www.nongnu.org/lzip/ |
| lzop | LZO | v1.04 | 2017 | https://www.lzop.org/ |
| lzturbo | LZ77 | v1.2 | 2014 | https://sites.google.com/site/powturbo/ |
| Nakamichi | LZSS | v1.0 | 2020 | http://www.sanmayce.com/Nakamichi/ |
| ppm | Prediction by partial matching | v1.0 | 2020 | https://github.com/rene-puschinger/ppm |
| qzip | Quicklz (https://github.com/robottwo/quicklz) library which is based on LZRW | v0.2 | 2011 | https://github.com/robottwo/quicklz |
| rans static | Arithmetic encoding | v1.0 | 2016 | https://github.com/jkbonfield/rans_static |
| rzip | Encodes large chunks of duplicated data then uses bzip2 | v1.0.6 | 2010 | https://rzip.samba.org/ |
| snzip | Snappy (https://github.com/google/snappy) which is a Google library based on ideas from LZ77 | v1.0.4 | 2016 | https://github.com/kubo/snzip |
| srank | Move-to-front encoding (symbol ranking) | v1.0 | 1997 | https://www.cs.auckland.ac.nz/~peter-f/FTPfiles/srank.c |
| xz | LZ77 with huge dictionary sizes and range encoding | v5.2.2 | 2015 | https://tukaani.org/xz/format.html |
| zlib | LZ77 | v1.2.11 | 2017 | https://github.com/madler/zlib |
| zip | LZ77 | v3.0 | 2008 | https://centos.pkgs.org/7/centos-x86_64/zip-3.0-11.el7.x86_64.rpm.html |
| zpipe | ZLib implementation of gzip and deflate | v1.0 | 2017 | https://github.com/skyformat99/zpipe |
| zstd | Facebook tool uses LZ77, finite-state-entropy, and Huffman encoding | v1.4.5 | 2020 | https://github.com/facebook/zstd |

**Table S1**: Most common general-purpose compression tools. Latest version for each tool was installed except for lrzip (the reason for this is provided in Supplementary Materials and Methods).

| **Category** | **Organism** | **Accession** | **Size (byte)** |
|---|---|---|---|
| Virus | Gordoniaphage GAL1 [1] | GCF 001884535.1 | 50,654 |
| Bacteria | WS1 bacterium JGI 0000059-K21 [2] | GCA 000398605.1 | 521,951 |
| Protist | Astrammina rara [2] | GCA 000211355.2 | 1,712,167 |
| Fungus | Nosema ceranae [2] | GCA 000988165.1 | 5,809,207 |
| Protist | Cryptosporidium parvumIowa II [2] | GCA 000165345.1 | 9,216,802 |
| Protist | Spironucleus salmonicida [2] | GCA 000497125.1 | 13,142,503 |
| Protist | Tieghemostelium lacteum [2] | GCA 001606155.1 | 23,672,980 |

| | | | |
|---|---|---|---|
| Fungus | Fusarium graminearumPH-1 [1] | GCF 000240135.3 | 36,915,673 |
| Protist | Salpingoeca rosetta [2] | GCA 000188695.1 | 56,150,373 |
| Algae | Chondrus crispus [2] | GCA 000350225.2 | 106,387,446 |
| Algae | Kappaphycus alvarezii [2] | GCA 002205965.2 | 341,012,624 |
| Animal | Strongylocentrotus purpuratus [1] | GCF 000002235.4 | 1,007,867,539 |
| Plant | Picea abies [2] | GCA 900067695.1 | 13,409,043,938 |

**Table S2.** Genome sequence datasets

| Dataset | No. of sequences | Size (byte) | Source | Date |
|---|---|---|---|---|
| Mitochondrion *[1]* | 9,402 | 245,282,526 | RefSeq ftp: ftp://ftp.ncbi.nlm.nih.gov/refseq/release/mitochondrion/mitochondrion.1.1.genomic.fna.gz ftp://ftp.ncbi.nlm.nih.gov/refseq/release/mitochondrion/mitochondrion.2.1.genomic.fna.gz | 15 March 2019 |
| NCBI Virus Complete Nucleotide Human *[3]* | 36,745 | 481,767,318 | NCBI Virus: https://www.ncbi.nlm.nih.gov/labs/virus/vssi/ | 11 May 2020 |
| Influenza [4] | 700,001 | 1,215,166,594 | Influenza Virus Database: ftp://ftp.ncbi.nih.gov/genomes/INFLUENZA/influenza.fna.gz | 27 April 2019 |
| Helicobacter [1] | 108,292 | 2,756,472,239 | NCBI Assembly: https://www.ncbi.nlm.nih.gov/assembly | 24 April 2019 |

**Table S3.** Other DNA datasets

| Genome | Size of genome in fasta format (byte) | Size after applying scripts (one-line format) (byte) |
|---|---|---|
| **DNA-Genome-Picea-abies-GCA_900067695.1-2016-11-09** | 13,409,043,938 | 11,960,690,255 |

| | | |
|---|---|---|
| **DNA-Helicobacter-2019-04-24** | 2,756,472,239 | 2,711,601,413 |
| **DNA-Influenza-2019-04-27** | 1,215,166,594 | 1,106,180,040 |
| **DNA-Genome-Strongylocentrotus-purpuratus-GCF_000002235.4-2015-03-10** | 1,007,867,539 | 659,082,267 |
| **DNA-NCBI-Virus-Complete-Nucleotide-Human-2020-05-11** | 481,767,318 | 470,877,038 |
| **DNA-Genome-Kappaphycus-alvarezii-GCA_002205965.2-2018-03-09** | 341,012,624 | 249,541,865 |
| **DNA-Mitochondrion-2019-03-15** | 245,282,526 | 241,606,731 |
| **DNA-Genome-Chondrus-crispus-GCA_000350225.2-2013-05-22** | 106,387,446 | 77,592,685 |
| **DNA-Genome-Salpingoeca-rosetta-GCA_000188695.1-2011-02-10** | 56,150,373 | 37,299,580 |
| **DNA-Genome-Fusarium-graminearum-PH-1-GCF_000240135.3-2008-11-21** | 36,915,673 | 34,133,023 |
| **DNA-Genome-Tieghemostelium-lacteum-GCA_001606155.1-2016-04-04** | 23,672,980 | 17,671,254 |
| **DNA-Genome-Spironucleus-salmonicida-GCA_000497125.1-2013-11-19** | 13,142,503 | 9,934,367 |
| **DNA-Genome-Cryptosporidium-parvum-Iowa-II-GCA_000165345.1-2007-02-26** | 9,216,802 | 7,012,512 |
| **DNA-Genome-Nosema-ceranae-GCA_000988165.1-2015-05-05** | 5,809,207 | 3,541,094 |
| **DNA-Genome-Astrammina-rara-GCA_000211355.2-2011-04-27** | 1,712,167 | 1,362,918 |
| **DNA-Genome-WS1-bacterium-JGI-0000059-K21-GCA_000398605.1-2013-05-16** | 521,951 | 509,551 |
| **DNA-Genome-Gordonia-phage-GAL1-GCF_001884535.1-2016-11-15** | 50,654 | 49,979 |
| **Total size in Megabytes** | **18,797.10** | **16,773.88** |

**Table S4.** Size of each genome before and after applying the script

| Tool | Compression command | Decompression command |
|---|---|---|
| **bcm** | bcm $genome_name | bcm -d -f $genome_name.bcm |
| **blzpack** | blzpack $genome_file $genome_name.blz | blzpack $genome_name.blz $genome_file |
| **brotli** | brotli $genome_file -c > $genome_name.br | brotli -d $genome_name.br -c > $genome_file |

| | | |
|---|---|---|
| **bsc** | bsc e $genome_file $genome_name.bsc -t -T | bsc d $genome_name.bsc $genome_file |
| **bzip2** | bzip2 -k -c $genome_file > $genome_name.bz2 | bunzip2 $genome_name.bz2 |
| **cmix** | cmix -c $genome_file ./$genome_name.cmix | cmix -d ./$genome_name.cmix ./$genome_file |
| **compress** | compress $genome_file -c > $genome_name.Z | compress -d $genome_name.Z -c > $genome_file |
| **freeze** | freeze -c $genome_file > $genome_name.F | freeze -d -c $genome_file > $genome_file |
| **gzip** | gzip $genome_file -c > $genome_name.gz | gunzip -d $genome_name.gz -c > $genome_file |
| **HE** | huffman -e $genome_name > $genome_name.hf | huffman -d $genome_name.hf > $genome_file |
| **hook** | hook08e c 256 2 1 255 $genome_file $genome_name.hk | hook08e d $genome_name.hk |
| **lizard** | lizard -f -c $genome_file > $genome_name.lzd | lizard -f -c -d $genome_name.lzd > $genome_file |
| **lrzip** | lrzip -q -N 1 -f $genome_file -o $genome_name.lrz | lrzip -d -q -N 1 -f $genome_name.lrz -o $genome_file |
| **lz4** | lz4 $genome_file -c > $genome_name.lz4 | lz4 -d $genome_name.lz4 -c > $genome_file |
| **lzb** | lzb < $genome_file > $genome_name.lzb | lzb -d < $genome_name.lzb > $genome_file |
| **lzfse** | lzfse -encode -i $genome_file > $genome_name.lzfse | lzfse -decode -i $genome_name.lzfse > $genome_file |
| **lzip** | lzip -c $genome_file > $genome_name.lz | lzip -d -c $genome_name.lz > $genome_file |
| **lzop** | lzop $genome_file -c > $genome_name.lzo | lzop -d $genome_name.lzo -c > $genome_file |
| **LzTurbo** | lzturbo -32 -o -f $genome_file > $genome_name.lzt | lzturbo -32 -o -f -d $genome_name.lzt > $genome_file |
| **nakamichi** | nakamichi $genome_file $genome_name.Nakamichi | nakamichi $genome_name.Nakamichi > $genome_file |
| **ppm** | ppm c $genome_file $genome_name.ppm | ppm d $genome_name.ppm $genome_file |
| **qzip** | qzip $genome_name | qunzip -d $genome_name.qz |
| **rans** | rANS_static -o1 $genome_file $genome_name.rans1 | rANS_static -d $genome_name.rans1 $genome_file |

| | | |
|---|---|---|
| **rzip** | rzip -k $genome_file -o $genome_name.rz | rzip -d -f $genome_name.rz |
| **snzip** | snzip -c $genome_file > $genome_name.sz | snzip -d -c $genome_name.sz > $genome_file |
| **srank** | srank $genome_file | srank $genome_name.sr |
| **xz** | xz -z -c $genome_file > $genome_name.xz | xz -d $genome_name.xz -c > $genome_file |
| **zip** | zip $genome_name.zip $genome_file | unzip $genome_name.zip |
| **zlib** | openssl zlib -e -in $genome_file > $genome_name.zlib | openssl zlib -d -in $genome_name.zlib > $genome_file |
| **zpipe** | zpipe < $genome_file > $genome_name.z | zpipe -d < $genome_name.z > $genome_file |
| **zstd** | zstd -f $genome_file -o $genome_name.zst | zstd -d -f $genome_name.zst |

**Table S5**. Commands used to compress files for each tool

### Further Analysis and Investigations of the Proposed Algorithm

Data binning is the underlying process in the proposed algorithm. Each sequence of length *w* (or of variable length) in *S* must be labeled and assigned to a bin with the corresponding label. Sequences in the same bin, which are assumed to be similar to each other, are then encoded and compressed together. Bins with different labels can be encoded and compressed using either a single encoding or a specific algorithm. The process used to compress a bin can be determined by the label and/or content of the bin.

This approach raises several questions that need to be addressed beyond this study. Given a set of concatenated or non-concatenated sequences sharing similar characteristics (e.g., the same Huffman tree), can an efficient or optimal encoding/compression solution be identified? What is the optimal classification/labeling method for bins? What is the optimal number of bins to be used? Is it better to use a window length that is fixed, variable, or dependent on the text/genome length?

Data binning has been used to solve compression problems for sequence reads, but not for genomic sequences or textual data. Assembltrie [5], BEETL [6], BdBG [7] FaStore [8], HARC [9], Mince [10], Orcom [11], and SCALCE [12] apply data binning to the compression of short variable-length sequences. There are several differences between these techniques and OST. First, these algorithms are typically applied to short sequences (of a few hundred bases in length), unlike genome or textual data. Second, the sequencing of such reads is typically conducted at high coverage. Hence, redundancy often exists between the reads, which is a motivating factor for the application of a binning process. Third, none of these algorithms preserve the order of the inputted reads.

The OST algorithm is designed primarily for long sequences in which there is no assumption of high repetitiveness in the input, as not all genomes or textual data exhibit such patterns and are potentially random. A key step in OST is to encode and record the order of the subsequences by recording the order of labels in $L$ to preserve the sub-sequence placement in the original input. This allows the data to be completely and exactly restored during the decompression process.
